# Exploration of the link between COVID-19 and gastric cancer from the perspective of bioinformatics and systems biology

**DOI:** 10.1101/2024.04.03.587916

**Authors:** Xiao Ma, Tengda Huang, Xiaoquan Li, Xinyi Zhou, Hongyuan Pan, Ao Du, Yong Zeng, Kefei Yuan, Zhen Wang

## Abstract

**Background:** Coronavirus disease 2019 (COVID-19), an infectious disease caused by severe acute respiratory syndrome coronavirus-2 (SARS-CoV-2), has caused a global pandemic. Gastric cancer (GC) poses a great threat to people’s health, which is a high- risk factor for COVID-19. Previous studies have found some associations between GC and COVID-19, whereas the underlying molecular mechanisms are not well understood.

**Methods:** We used a bioinformatics and systems biology approach to investigate the relationship between GC and COVID-19. The gene expression profiles of COVID-19 (GSE196822) and GC (GSE179252) were downloaded from the Gene Expression Omnibus (GEO) database. After identifying the shared differentially expressed genes (DEGs) for GC and COVID-19, functional annotation, protein-protein interaction (PPI) network, hub genes, transcriptional regulatory networks and candidate drugs were analyzed.

**Results:** A total of 209 shared DEGs were identified to explore the linkages between COVID-19 and GC. Functional analyses showed that Immune-related pathway collectively participated in the development and progression of COVID-19 and GC. In addition, there are selected 10 hub genes including *CDK1*, *KIF20A*, *TPX2*, *UBE2C*, *HJURP*, *CENPA*, *PLK1*, *MKI67*, *IFI6*, and *IFIT2*. The transcription factor/gene and miRNA/gene interaction networks identified 38 transcription factors (TFs) and 234 miRNAs. More importantly, we identified ten potential therapeutic agents, including ciclopirox, resveratrol, etoposide, methotrexate, trifluridine, enterolactone, troglitazone, calcitriol, dasatinib and deferoxamine, some of which have been reported to improve and treat GC and COVID-19. This study also provides insight into the diseases most associated with mutual DEGs, which may provide new ideas for research on the treatment of COVID-19.

**Conclusions:** This research has the possibility to be contributed to effective therapeutic in COVID-19 and GC.

## Introduction

Coronavirus disease 2019 (COVID-19) has already caused a global pandemic in just a few years after its appearance, posing a significant health challenge worldwide, with the pandemic having killed more than 6 million people[1]. As the pandemic progresses, new variants of the severe acute respiratory syndrome coronavirus 2 (SARS-CoV-2) have emerged. The World Health Organization (WHO) has declared the variants of concern, including Alpha, Beta, Gamma, Delta, and Omicron[2,3]. SARS-CoV-2 contact with the angiotensin converting enzyme 2 (ACE2) receptor upon entry into the body and replicates and releases within the epithelium, thereby infecting surrounding cells[4]. As an important component of the renin-angiotensin (RAS) system, ACE2 has been learned as a membrane receptor for SARS-COV-2[5]. In addition, SARS-CoV-2 also enters host cells with the primary or auxiliary help of host proteases transmembrane protease serine 2 (TMPRSS2), FURIN[6], glucose-regulating protein 78 (GRP78) receptor[7], dipeptidyl peptidase 4 (DPP4)[8], cluster of differentiation 147 (CD147) transmembrane protein[9], tyrosine-protein kinase receptor UFO (AXL)[10], phosphatidylinositol 3-phosphate 5-kinase (PIKfyve)[11], two pore channel subtype 2 (TPC2)[12] and cathepsin L[13]. SARS-CoV-2 infection alters alveolar vascular permeability, which in turn leads to lung injury such as pulmonary edema, DIC, and pulmonary ischemia[14]. Not only that, SARS-CoV-2 can also spread to the brain, heart, gastrointestinal tract and other important organs through blood, which may cause adverse effects such as cerebral hemorrhage, coma, paralysis and eventually death[15–19]. Due to the genetic diversity and frequent genome reorganization of SARS-CoV-2, COVID-19 is likely to evolve and become seasonal epidemics[20]. Clinical and epidemiological data suggest that certain underlying diseases, such as cancer, hypertension, cardiovascular disease and diabetes, make the organism more susceptible to SARS-CoV-2 infection and may lead to more serious lung damage and even patient death[21]. How to treat COVID-19 infection in people with underlying diseases, including cancer, is of great research value and clinical significance.

Gastric cancer (GC) is a main worldwide health concern and is currently the fourth leading cause of cancer-related deaths worldwide[22]. The incidence of GC has a relatively strong geographic pattern, with the highest morbidity in East Asia, some South American and Eastern European countries, and the lowest incidence in Africa and North America. Globally, more than 70% of GC happens in developing countries. Patients with gastric cancer have poor prognosis and low long-term survival rates[23]. Some investigators have suggested that GC patients are more sensitive to SARS-CoV-2 infection and that immunotherapy and radiotherapy within 3 months of a diagnosis of COVID-19 are risk factors for death[24]. The presence of ACE2 receptors in the gastric mucosa indicates that these cells may also be infected by SARS-CoV-2[25]. In addition, a Mendelian randomization study revealed a suggestive causal relationship between SARS-CoV-2 infection and an increased risk of gastric cancer[26]. Therefore, finding the molecular mechanism of the interaction of COVID-19 on gastric cancer patients is likely to inform subsequent studies or treatments.

In this study, transcriptome profiles were obtained from the National Center for Biological Information (NCBI)-Gene Expression Omnibus (GEO) database. The datasets of COVID-19 and GC were studied to find differentially expressed genes (DEGs) for both diseases. These sets of DEGs were then compared to gain mutual DEGs. Moreover, the biological function of the common DEGs was analysed to gain insights into its impact on disease onset and progression. A protein-protein interaction (PPI) network was used to identify hub genes with the most obvious interactions and narrow down potential biomolecules. Next, the hub genes were used to establish the gene-regulatory network, predict potential drugs, and complete the gene-disease association network. Figure 1 shows the successive workflow of this study. This study will identify the shared pathogenesis between gastric cancer and COVID-19 and analyse possible drug candidates, potentially providing new ideas to assist in the treatment of GC and COVID-19.

**Figure 1.**
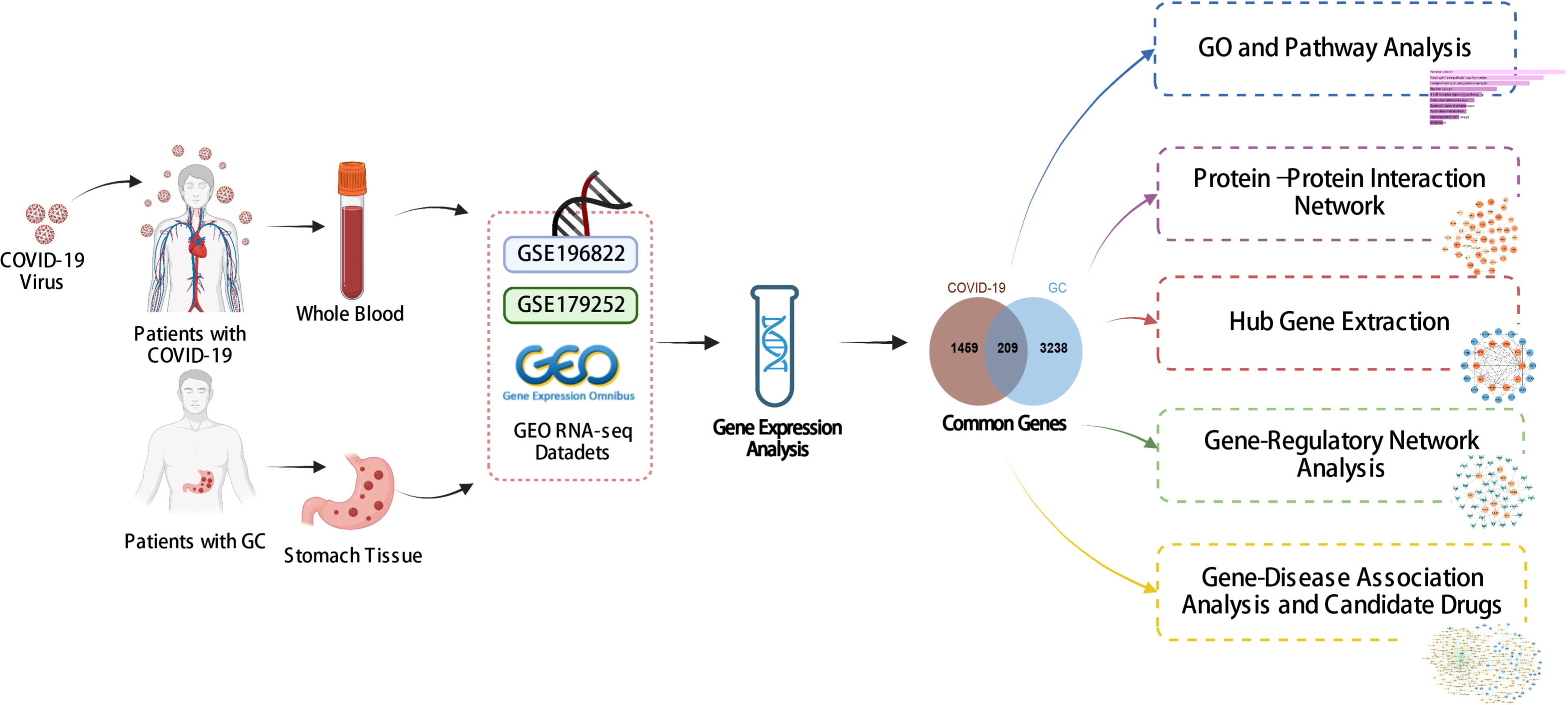
The overall work of this study.

## Materials and methods

### Data source

To determine the mutual genetic interrelationship between SARS-CoV-2 infection and gastric cancer, we used the GEO database of NCBI to obtain RNA-seq datasets. For SARS-CoV-2 patients, we used GSE196822 dataset[27], which includes whole-blood transcriptome profiling of 34 COVID-19 patients and 9 healthy controls. The data came from high-throughput sequencing using the Illumina HiSeq 4000 (Homo sapiens). Gastric cancer (GSE179252)[28] consists of 38 gastric tumors and paired normal 38 gastric tissues which was based on Illumina HiSeq 4000 (Homo sapiens).

### Identification of DEGs and common DEGs between COVID-19 and gastric cancer

The key target of the analysis is to find the DEGs for the datasets GSE196822 and GSE179252. The DEseq2 package[29] of R software (version 4.2.0) was used to identify the DEGs with false-discovery rate (FDR) < 0.05 and |log2 Fold Change| > 1.

To extract the shared DEGs between COVID-19 and gastric cancer, an online VENN analysis tool called Jvenn was used[30].

### Gene ontology and pathway enrichment analysis

The use of gene ontology (GO) enrichment methods is widespread for demonstrating the relationship between genes and GO terms, and the GO database is a comprehensive resource on gene and gene product functions[31]. GO annotated sources for biological process (BP), molecular function (MF), and cellular component (CC) were retrieved from the GO database[32]. To identify pathways shared by GC and COVID-19, we considered the following three repositories as the origin of pathway classification: WikiPathways, Reactome, and the Kyoto Encyclopedia of Genes and Genomes (KEGG). We used Enrichr[33] (https://maayanlab.cloud/Enrichr/) for gene ontology and pathway enrichment investigations. To quantify the top pathways and functional items, a standardized index with a P-value < 0.05 was utilized.

### Protein–Protein interaction (PPI) network analysis and hub genes extraction

STRING (an online tool https://string-db.org/, version 11.5) has been used to construct the PPI network with a filter condition (combined score>0.5)[34]. All the common DEGs between COVID-19 and GC were used to build the PPI network. Then, the PPI network was consumed into Cytoscape (v.3.9.1) for visual representation and hub genes’ recognition[35]. We used Maximal Clique Centrality (MCC) method of Cytohubba (a plugin of Cytoscape, http://apps.cytoscape.org/apps/cytohubba) to identify the top 10 hub genes from the PPI network[36]. At the same time, Cytohubba’s proximity ranking characteristics helped us to identify the shortest reachable pathways linking hub genes.

### Identification of miRNAs–gene and transcription factors–gene interactions

Transcription factors (TFs) are proteins attached to specific genes that control the genetic information’s transcription rate; as such, they are essential for molecular insight[37]. Our approach involved utilizing the NetworkAnalyst platform (version 3.0)[38] to identify topologically feasible TFs from the JASPAR database that could be potentially integrated with hub genes. JASPAR is a publicly accessible database that compiles information on TFs across six taxonomic groups for various species[39]. miRNAs targeting gene interactions are also included in studies to identify miRNAs that tend to bind to gene transcripts and thus negatively impact protein production. We used Network-Analyst to analyse miRNAs-gene interactions from Tarbase (version

8.0)[40] databases.

### Identification of drug candidates

Another emphasis of this study was to use the mutual DEGs of COVID-19 and GC to predict protein-drug interactions (PDIs) or drug molecule recognition. Using Enrichr’s disease/drug functions, based on Hub genes, drug molecules were predicted from the Drug Signatures Database (DSigDB, http://tanlab.ucdenver.edu/DSigDB)[41], which contains 17,389 unique chemicals that span 19,531 genes and has 22,527 gene sets.

### Gene-disease association analysis

DisGeNET (http://www.disgenet.org/)[42] is a platform that integrates and standardizes data on genes associated with diseases from diverse sources. Currently, DisGeNET has information on about 24,000 illnesses and features, 17,000 genes, and 117,000 genetic variations. To identify the relationship between relevant diseases and common DEGs, we use DisGeNET, Network-Analyst and Cytoscape to investigate the relationship between genes and diseases.

## Results

### Determination of DEGs and common DEGs of GC and COVID-19

To investigate the correlation and influence between GC and COVID-19, we analysed human RNA-seq datasets from GEO and identified shared DEGs that may trigger both COVID-19 and GC. In this study, 1,668 genes were found to be differentially expressed in COVID-19, including 839 up regulated DEGs and 829 down regulated DEGs (Supplemental file 1: Table S1). Similarly, 3,447 DEGs were identified in the GC data, including 1045 up regulated DEGs and 2402 down regulated DEGs (Supplemental file 2: Table S2). The information about the two datasets has been integrated in Table 1. To find shared DEGs between GC and COVID-19, we performed a cross-comparative evaluation using Jvenn and identified 209 common DEGs in both datasets (Figure 2 and Supplemental file 3: Tables S3). There are multiple genes in common between GC and COVID-19, suggesting some similarity between the two diseases.

**Figure 2.**
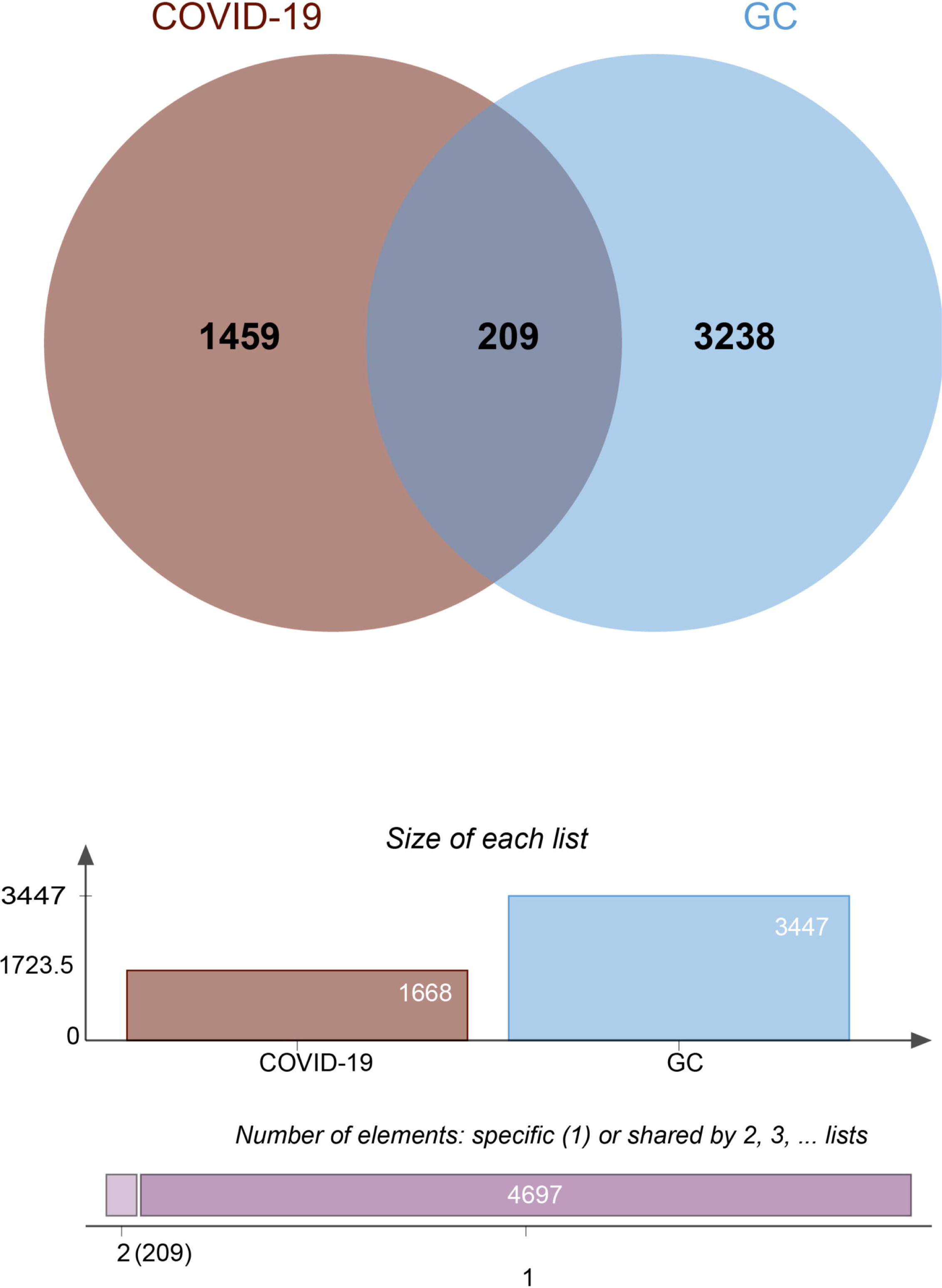
The Venn diagram showed 209 shared DEGs between COVID-19 and GC.

**Table 1.**
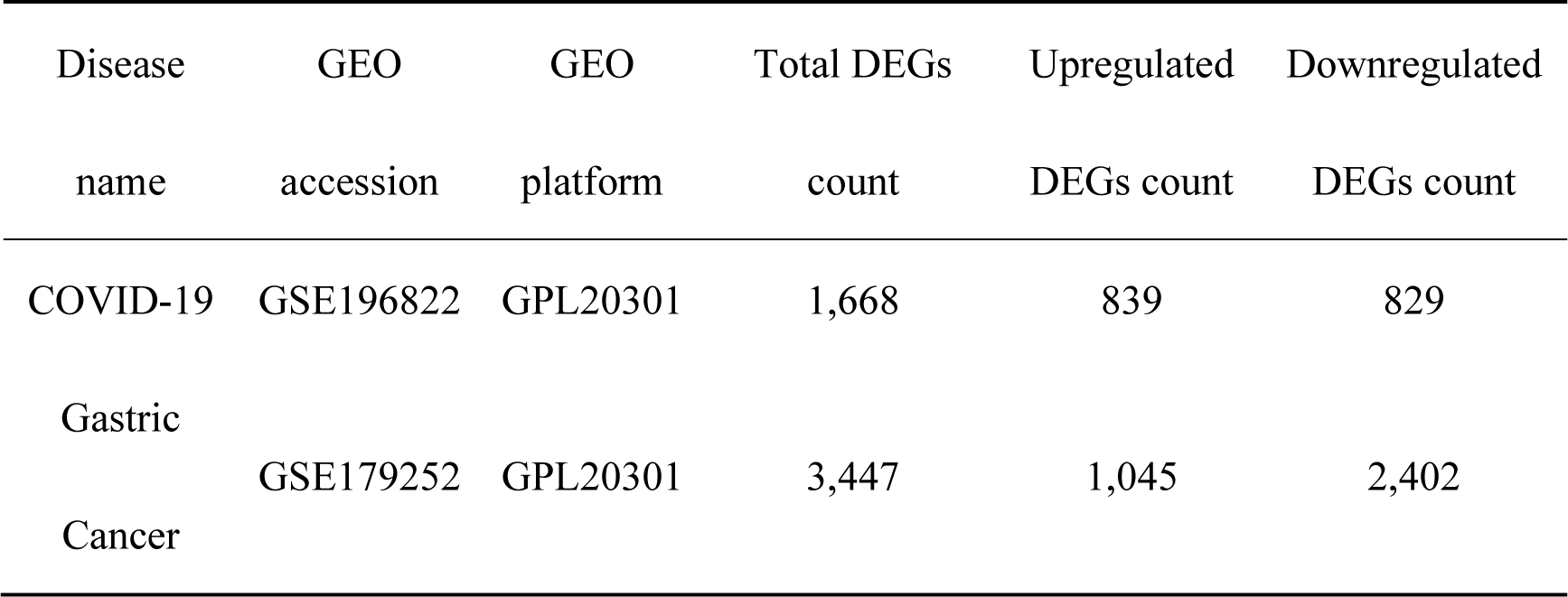
Overview of datasets with their geo-features and their quantitative measurements in this analysis

### Analyses of GO and pathway enrichment

To investigate the enrichment pathways and biological significance of common DEGs between COVID-19 and GC, we used Enrichr for gene functional annotation. Fig 3 and Table 2 summarized the top 10 enriched GO categories in the biological process, molecular function and cellular component categories. Notably, common DEGs were significantly enriched in immune-related pathways, which include neutrophil degranulation (GO:0043312), neutrophil activation involved in immune response (GO:0002283), neutrophil mediated immunity (GO:0002446), innate immune response (GO:0045087), defense response to virus (GO:0051607), receptor ligand activity (GO:0048018), and cytokine activity (GO:0005125).

**Table 2.**
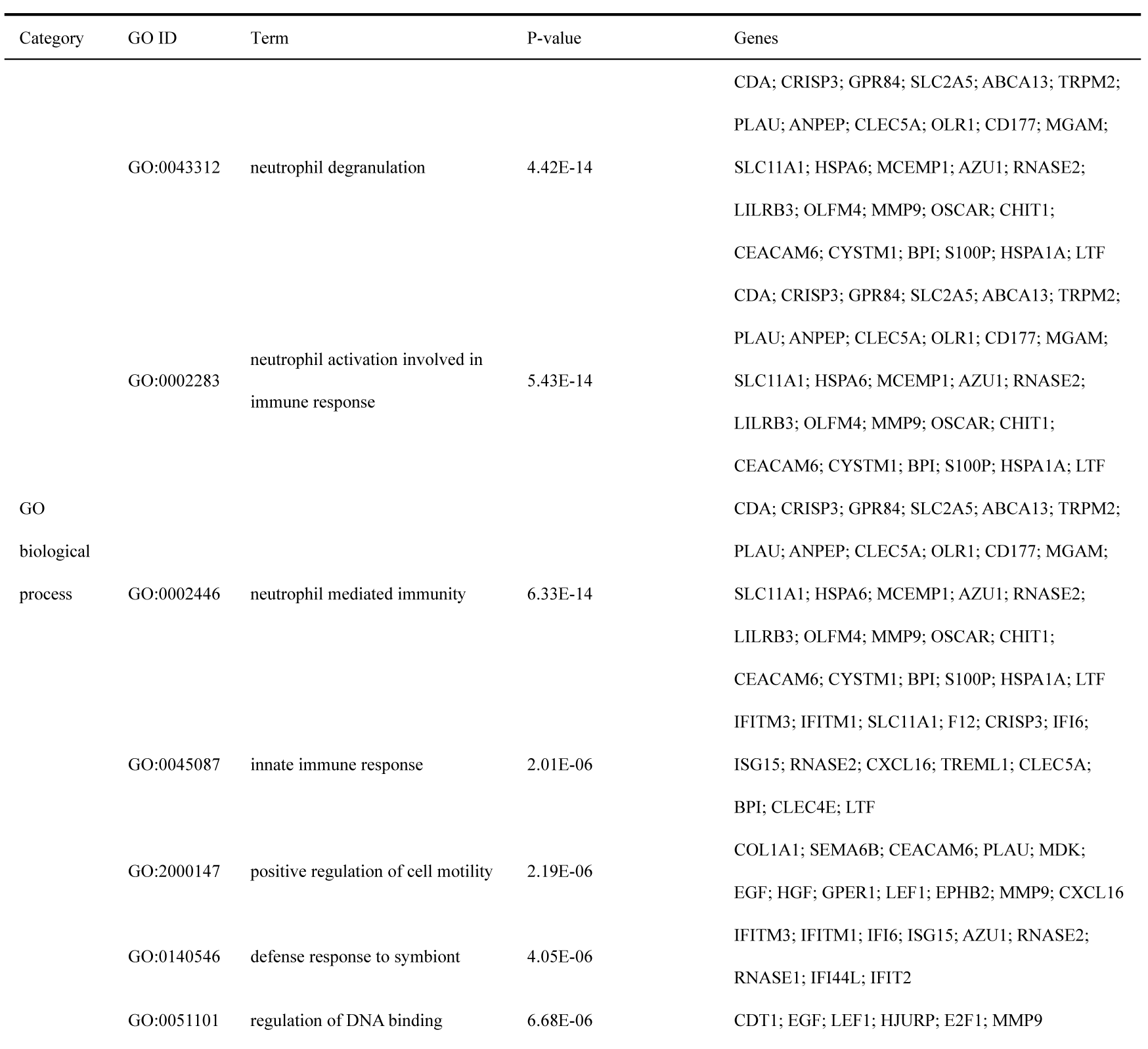
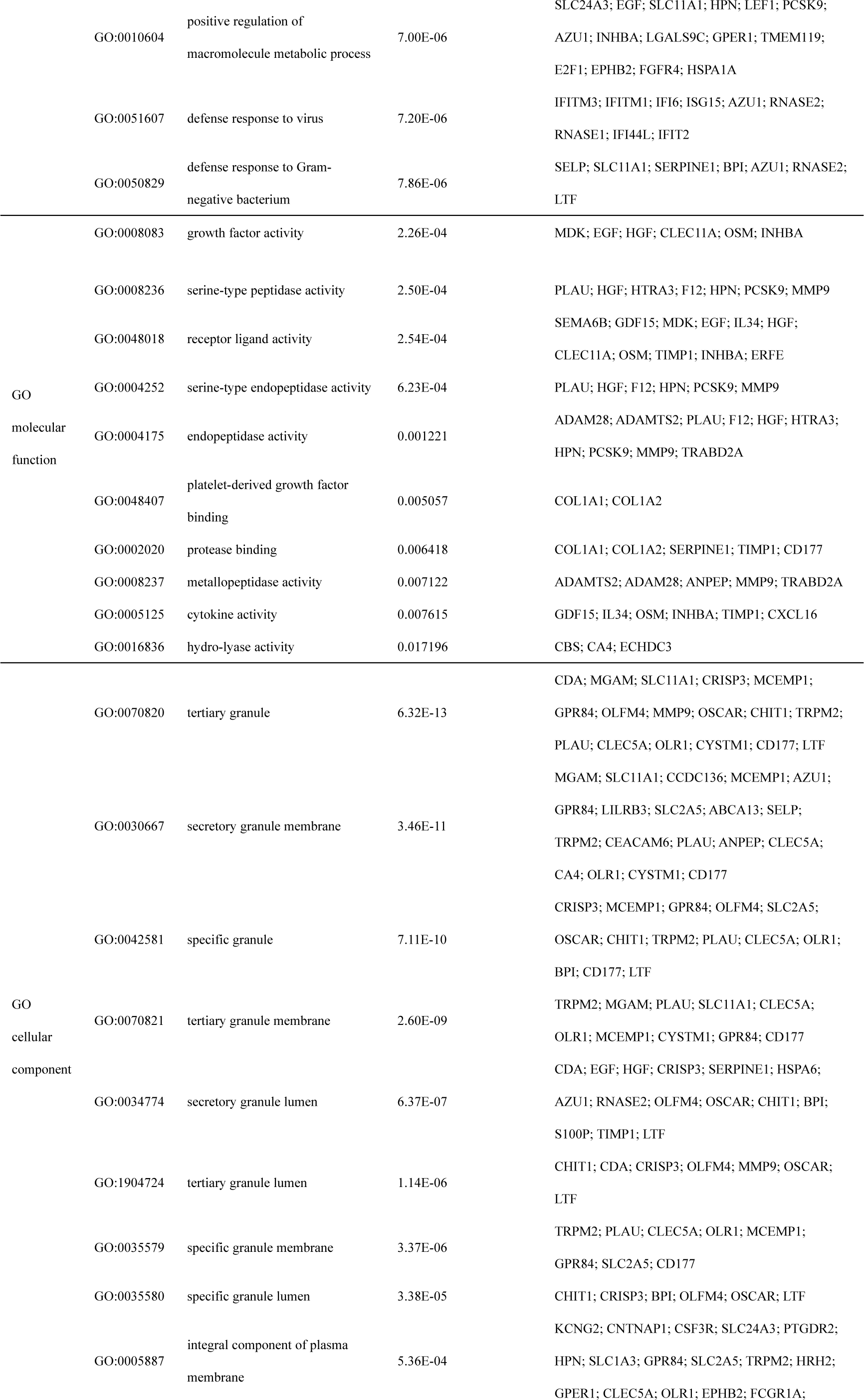
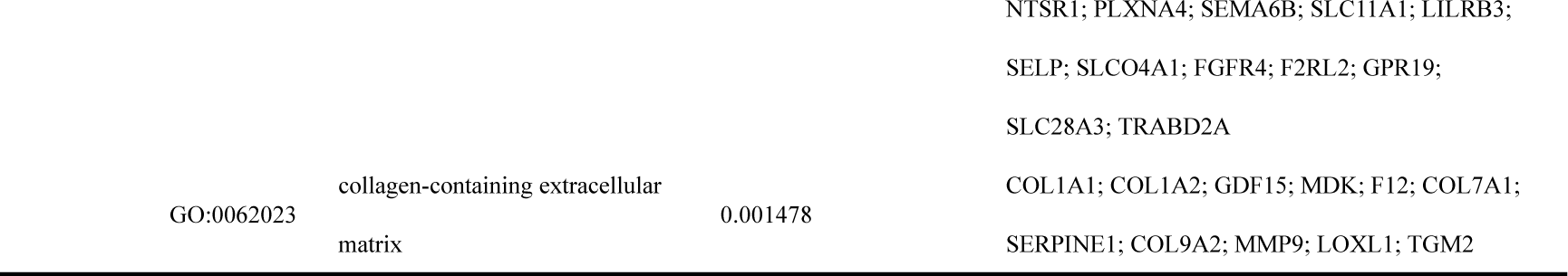
Ontological analysis of common DEGs between SARS-CoV-2 and GC

**Figure 3.**
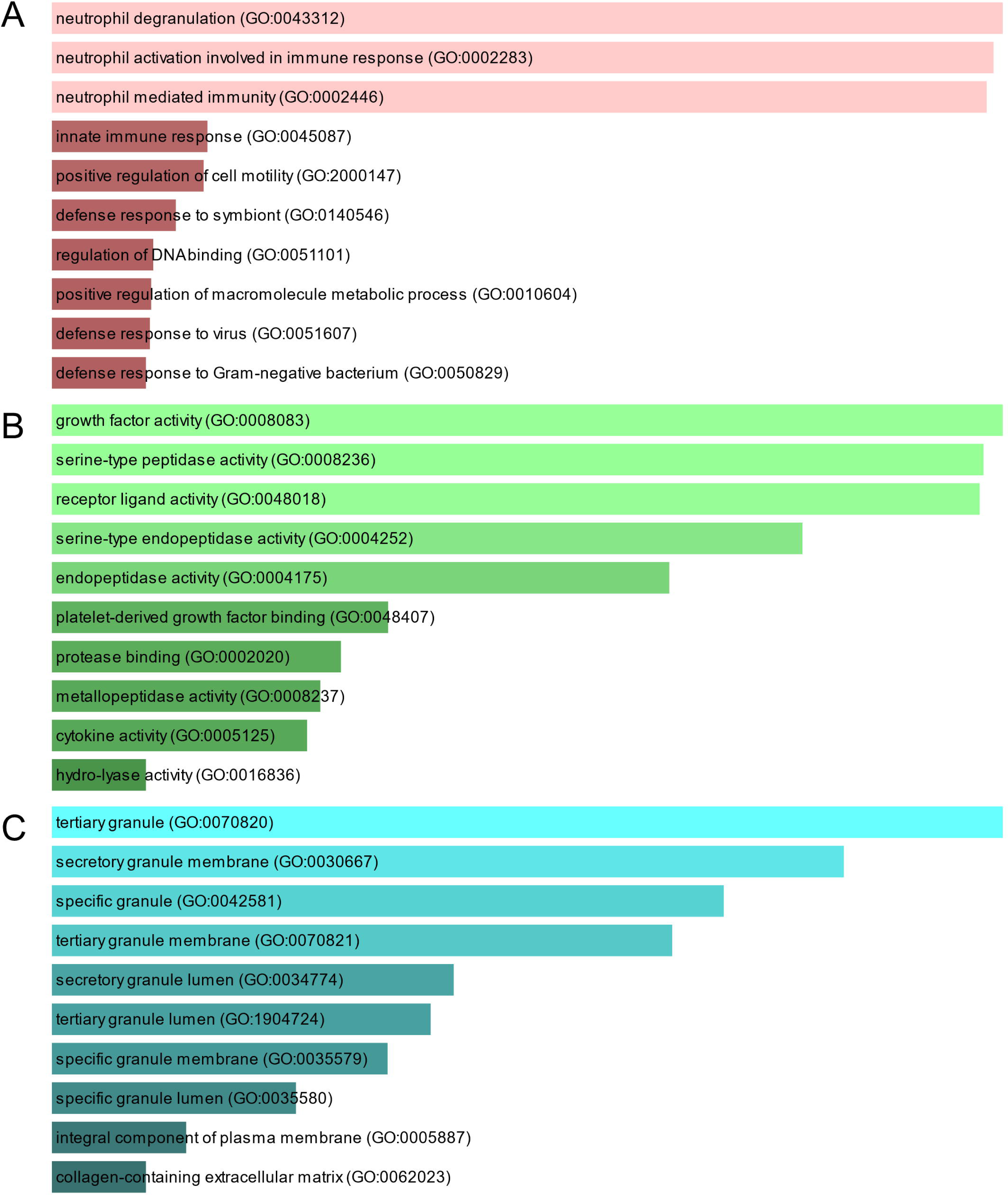
The bar graphs depicting the gene ontology enrichment analysis of shared DEGs between COVID-19 and gastric cancer. **A:** biological process. **B:** molecular function. **C:** cellular component. The lighter the color, the more significant it is.

Pathway analysis can highlight how underlying molecular and biological processes interact[43]. Figure 4 and Table 3 show the main pathways of common DEGs enrichment in WikiPathways, Reactome, and KEGG. Pathway enrichment analysis showed that common DEGs are mainly involved in the regulation of immune-related pathways, including TGF-beta Receptor Signaling WP560, Neutrophil Degranulation R-HSA-6798695, Immune System R-HSA-168256, Innate Immune System R-HSA- 168249, Immunoregulatory Interactions Between A Lymphoid And A non-Lymphoid Cell R-HSA-198933, Transcriptional Regulation Of Granulopoiesis R-HSA-9616222, Interferon Alpha/Beta Signaling R-HSA-909733 and B cell receptor signaling pathway. These results provide strong evidence that these common DEGs play a role in the onset and development of COVID-19 and GC through immune-related pathways.

**Figure 4.**
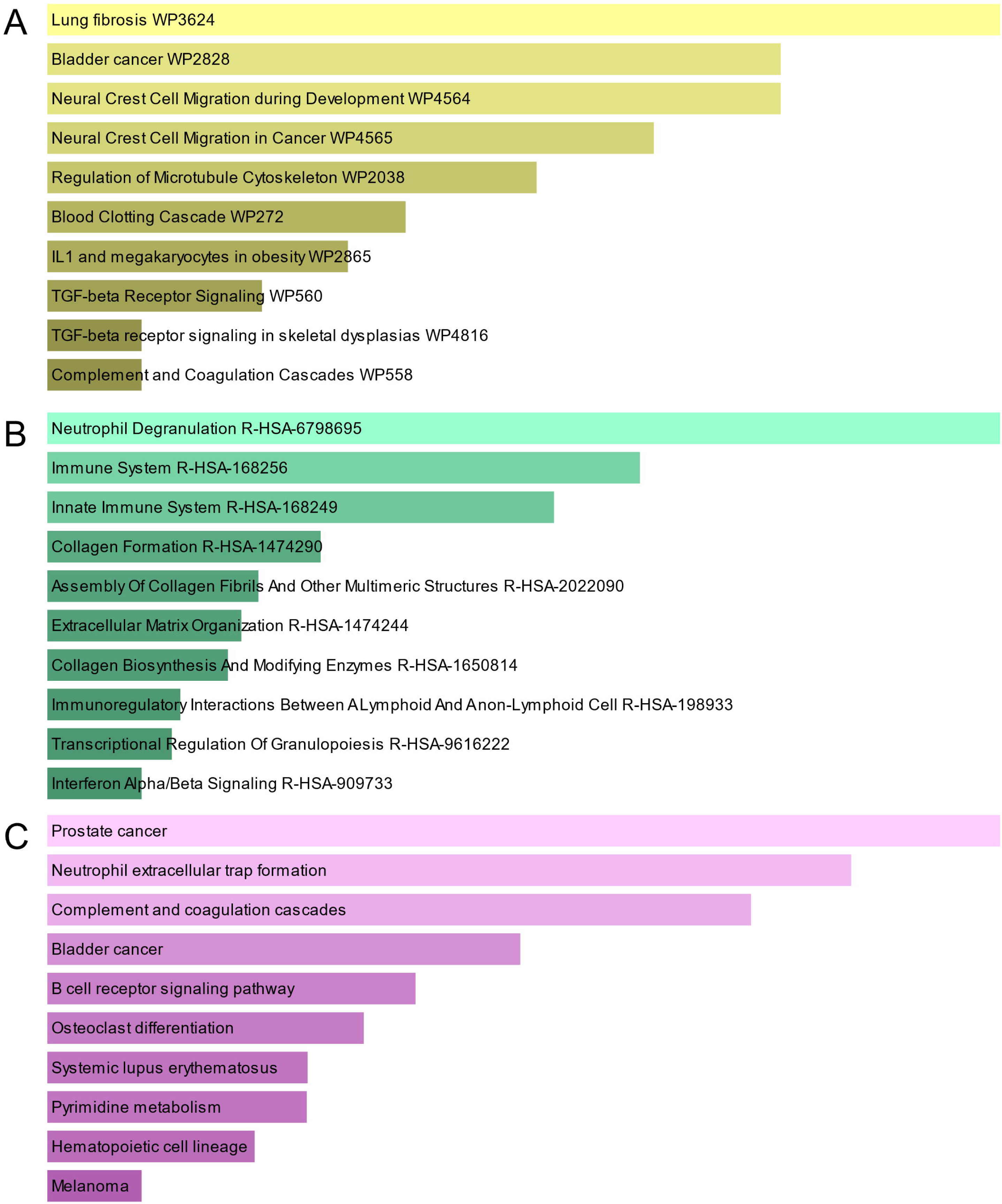
The bar graphs of pathway enrichment analysis of mutual DEGs between COVID-19 and GC. **A:** WikiPathway. **B:** Reactome pathway. **C:** KEGG. The lighter the color, the more significant it is.

**Table 3.**
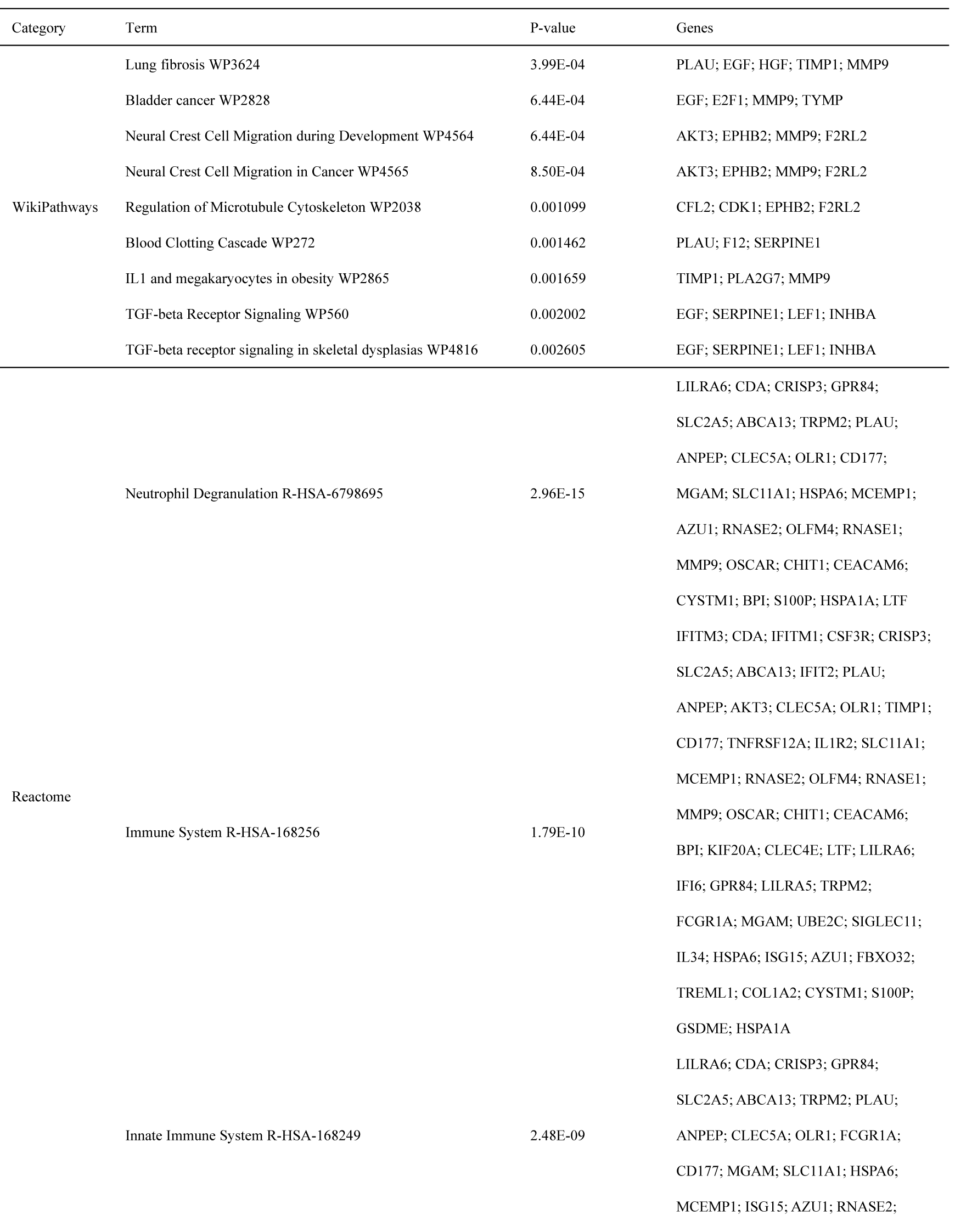
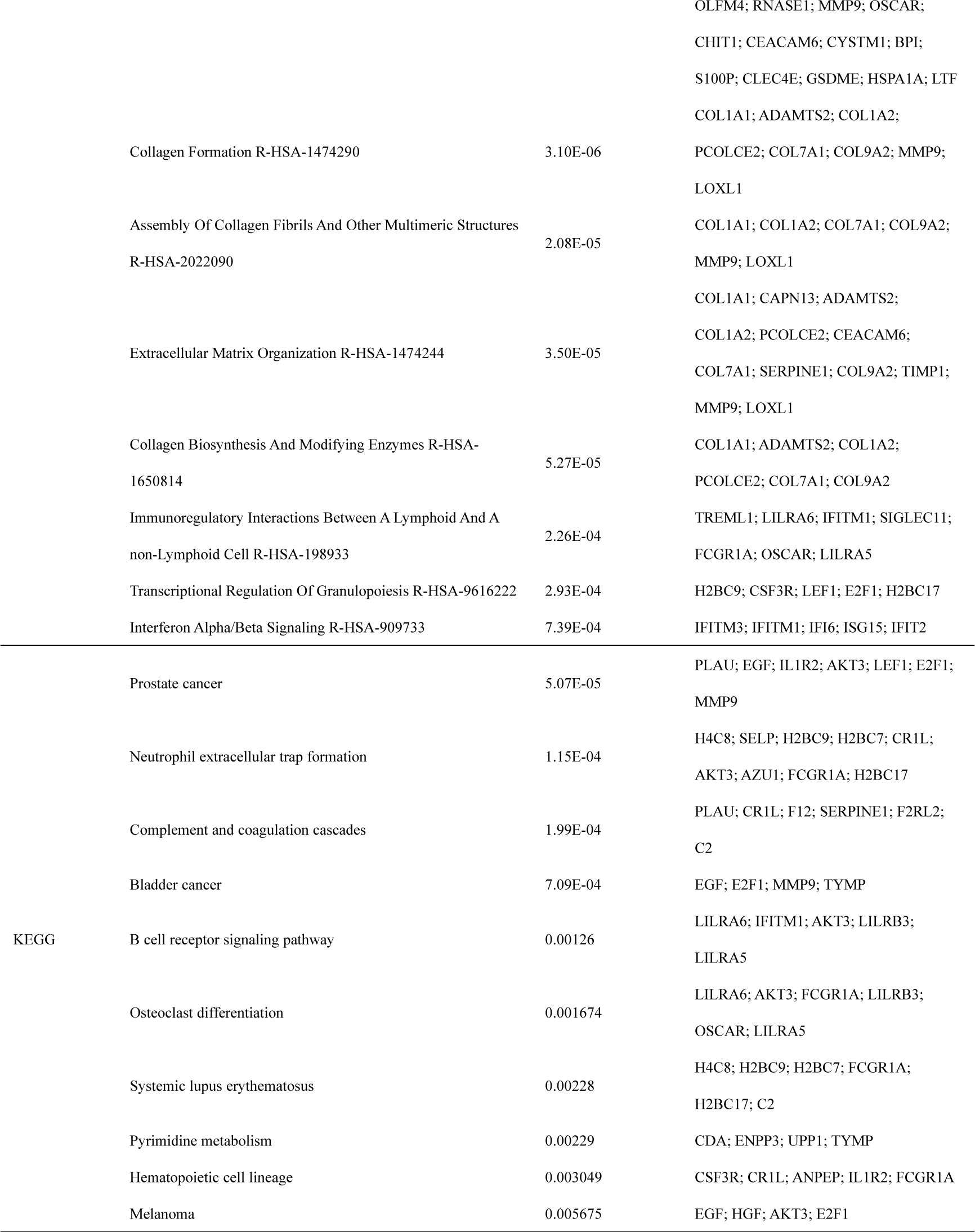
Pathway enrichment analysis of common DEGs between SARS-CoV-2 and GC

## Building PPI network and selecting of hub genes

PPI networks can visualize the interrelationships between different proteins and can help us understand the underlying mechanisms by which proteins interact[44]. Using STRING and Cytoscape, we built and visualized a PPI network of shared DEGs between COVID-19 and GC, which encompasses 46 nodes and 82 edges, as depicted in Figure 5. The most entangled nodes among them are hub genes. These hub genes have the potential to serve as biomarkers and may offer novel insights into therapeutic approaches. The top 10 hub genes with the highest MCC scores were identified using the cytoHubba plugin of Cytoscape, namely *CDK1*, *KIF20A*, *TPX2*, *UBE2C*, *HJURP*, *CENPA*, *PLK1*, *MKI67*, *IFI6*, and *IFIT2*(Figure 6 and Supplemental file 4: Tables S4). **Identifying of transcription regulatory network**

**Figure 5.**
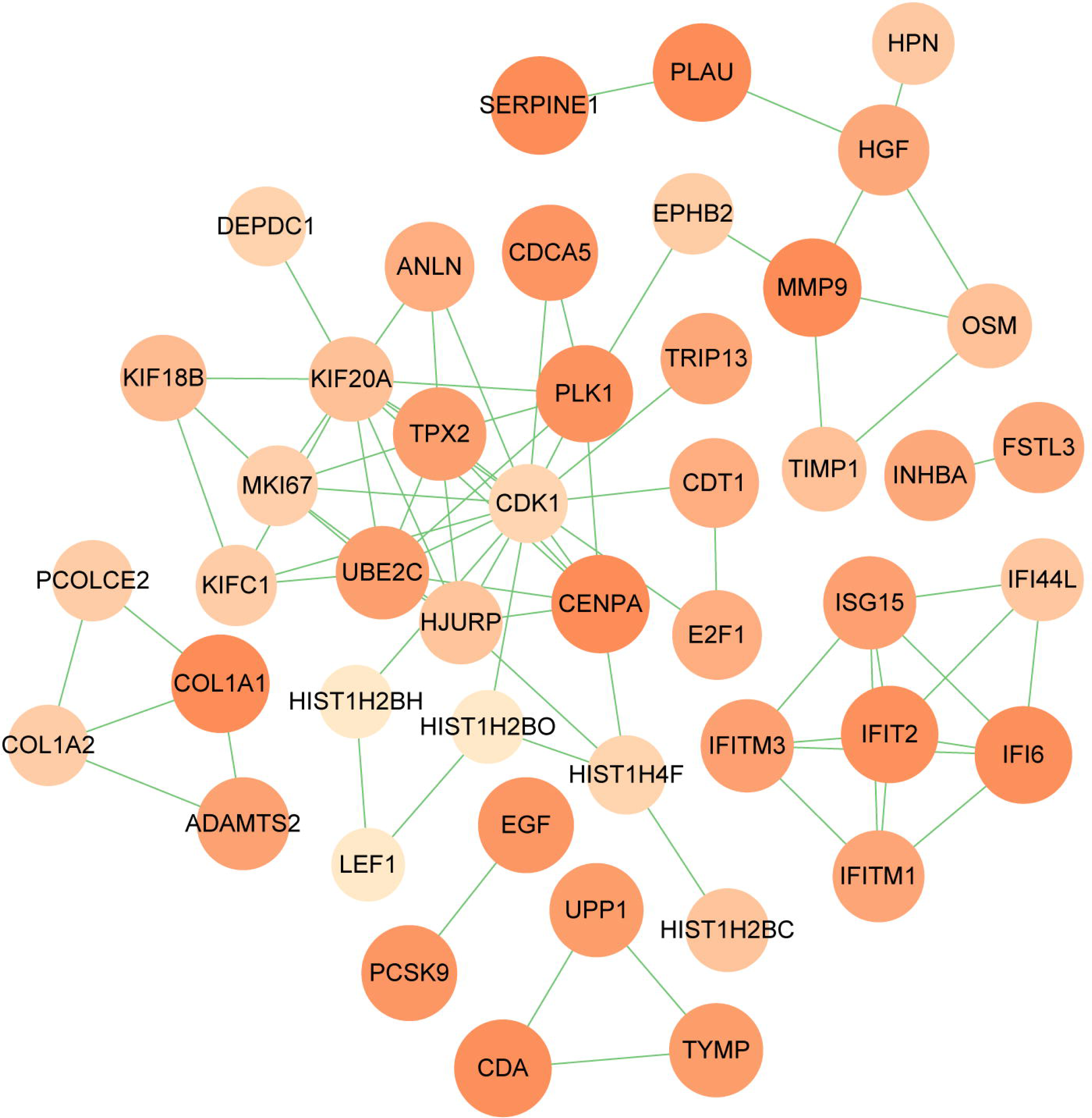
PPI network of shared DEGs between COVID-19 and GC. The DEGs in the figure are represented by circles and the edges mean the connections between the nodes. The PPI network consists of 46 nodes and 82 edges.

**Figure 6.**
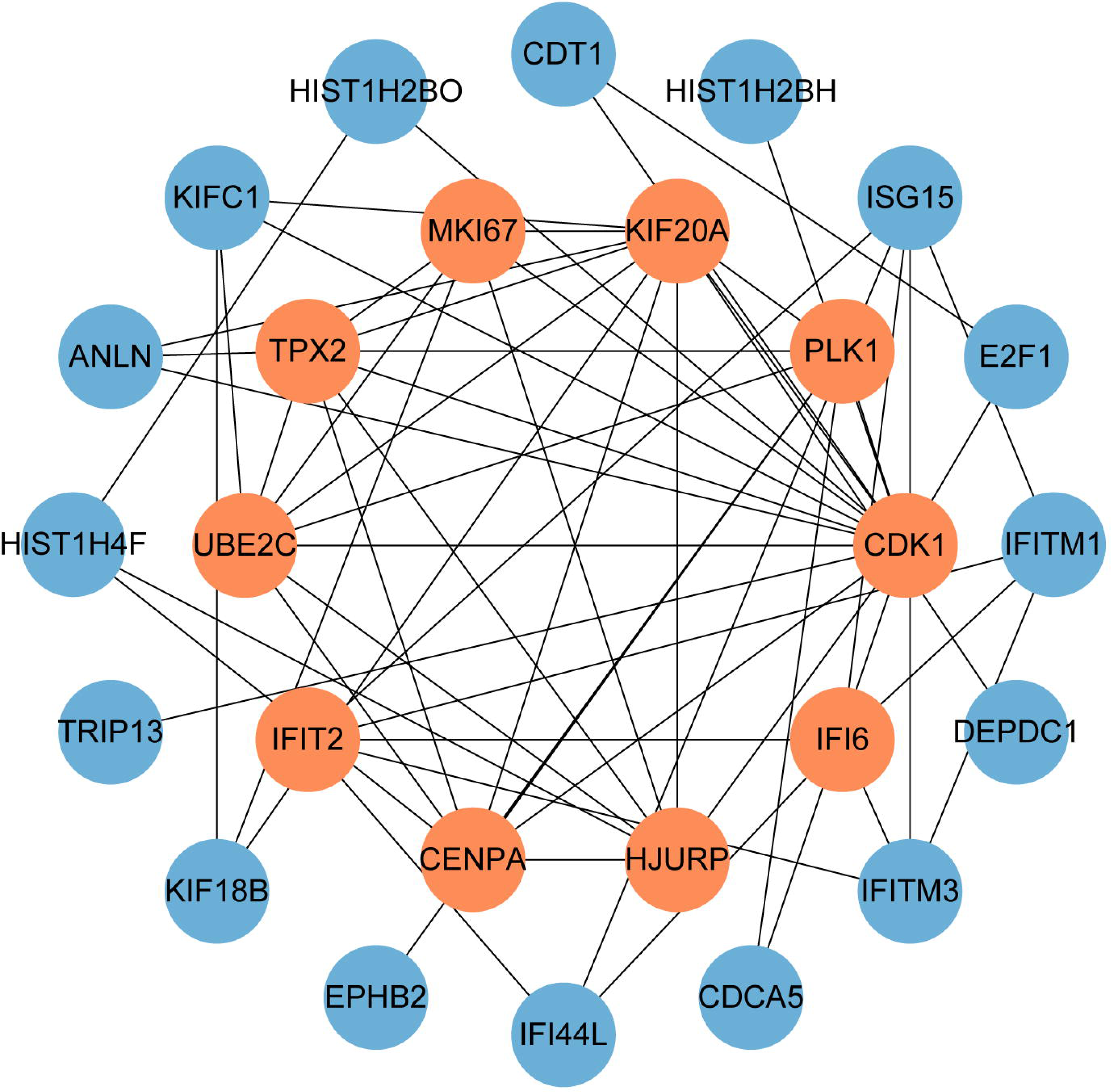
The hub genes were obtained from the PPI network. The nodes in orange represent the top 10 prominent hub genes and the interactions among them and other molecules. This network contains 26 nodes and 60 edges.

To figure out how hub genes modulate COVID-19 and GC at the transcriptional level, this study also investigated the interaction between TFs and genes, as well as miRNAs. In this study, a network-based approach was used to decode regulatory transcription factors and miRNAs as a way to gain insight into hub genes and find substantial changes that occur at the transcriptional level. As shown in Figure 7, the figure exhibits the interaction network of 38 transcription factors, such as ESR2, REL, SRY, SPIB, NR2F1, BRCA1, FOS, CREB1, FOXC1 and GATA2 (Supplemental file 5: Tables S5). Similarly, Figure 8 represents the interaction network of miRNAs regulators and hub genes, containing 234 miRNAs, such as hsa-mir-16-5p, hsa-mir-192-5p, hsa-mir-215-5p, hsa-mir-92a-3p, hsa-mir-193b-3p, hsa-let-7e-5p, hsa-mir-1283, hsa-mir-218-5p, hsa-mir-1- 3p and hsa-mir-671-5p (Supplemental file 6: Tables S6).

**Figure 7.**
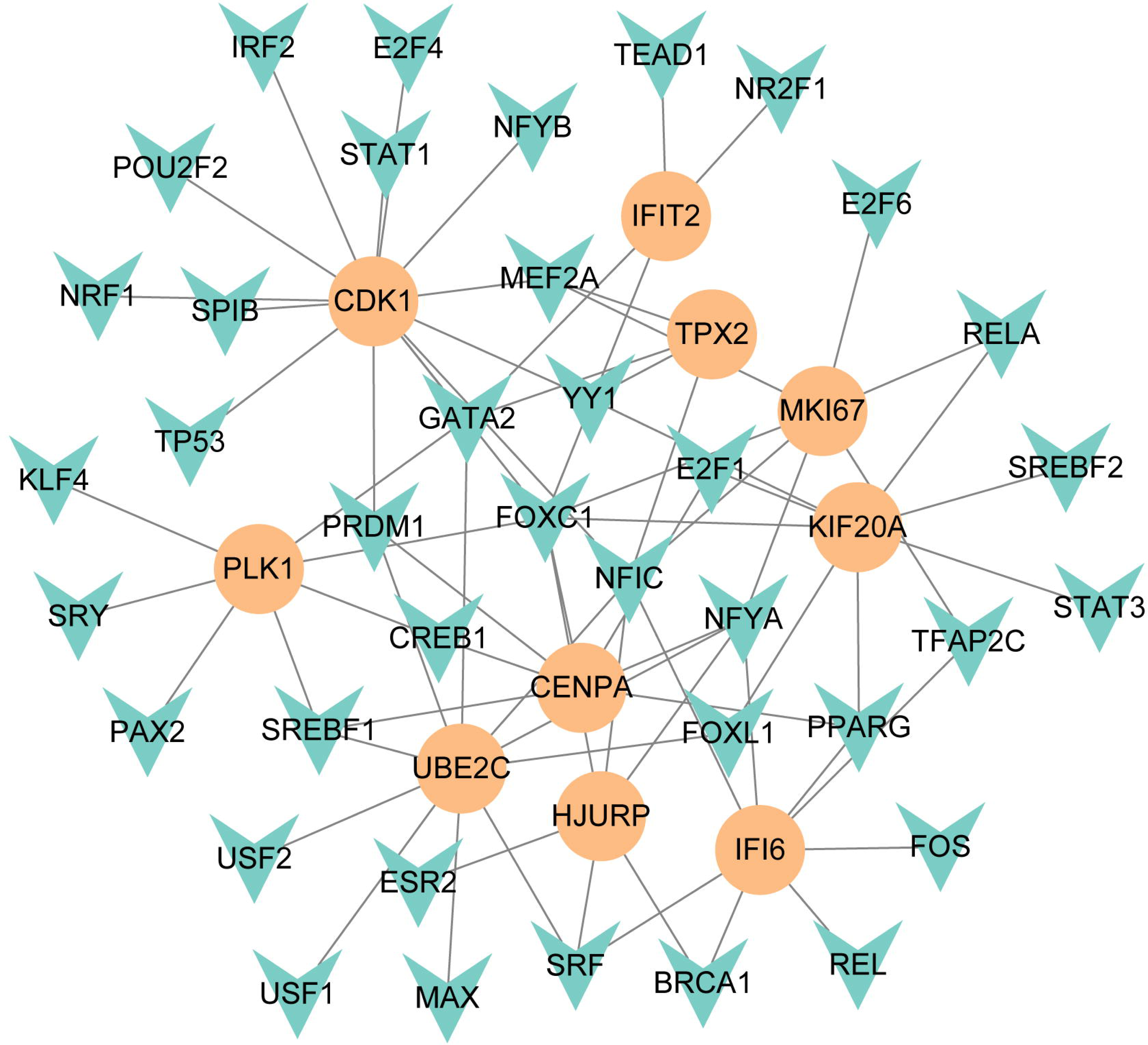
The regulatory network of TFs-gene. The circles represent hub gene, and transcription factor was represented by quadrilateral shape. The network contains 38 nodes and 75 edges.

**Figure 8:**
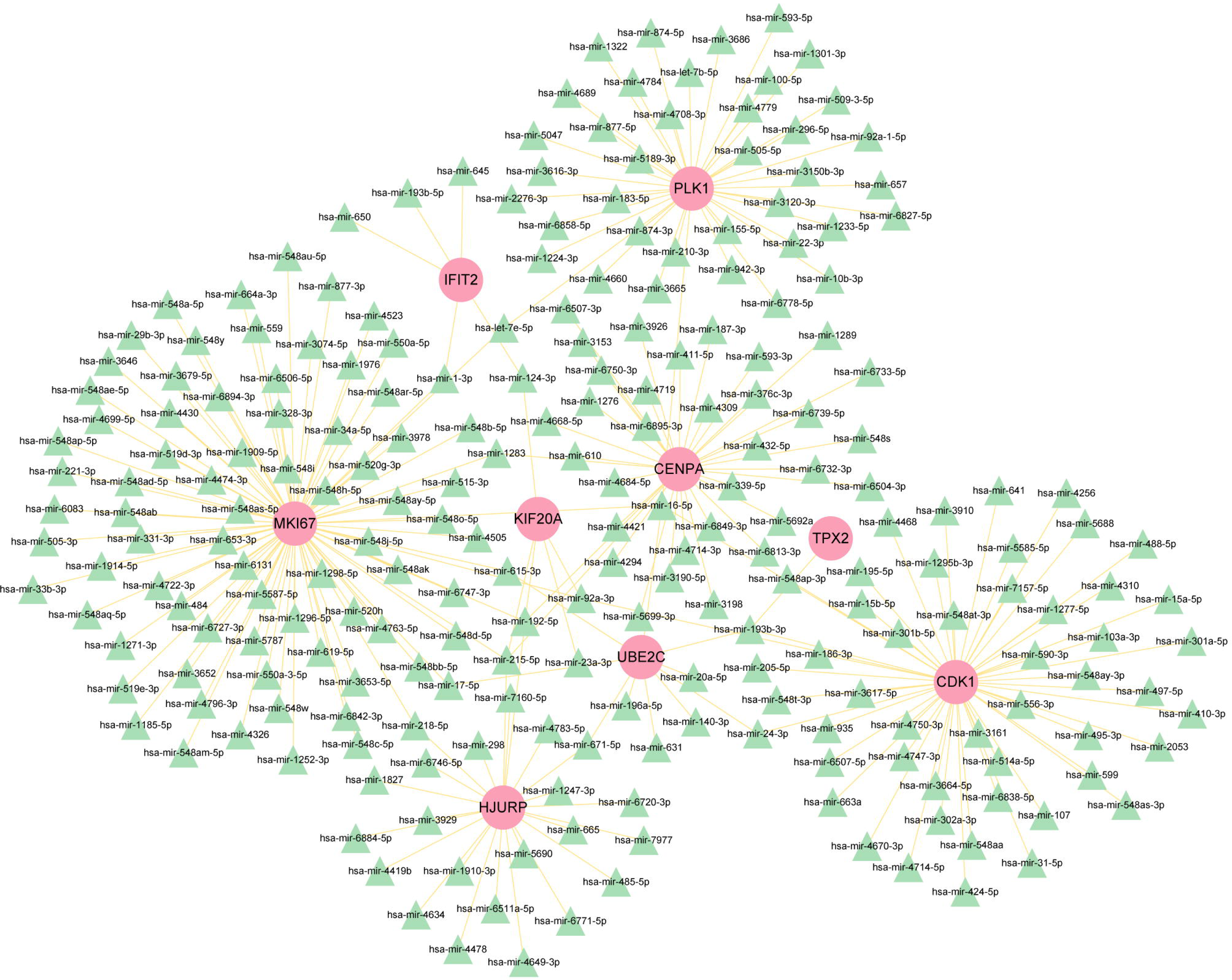
The regulatory network of miRNAs-gene. The pink circle represents the hub gene and the green triangle represents the miRNA. The network contains 243 nodes and 256 edges.

### Determination of candidate drugs

To search for potential drugs to treat COVID-19 and GC, possible drug molecules were predicted based on the transcriptional characteristics from the DSigDB database[45]. The top 8 compounds were identified according to their P-values (Table 4). The potential drug compounds were ciclopirox, resveratrol, etoposide, methotrexate, trifluridine, enterolactone, troglitazone, calcitriol, dasatinib and deferoxamine. These drugs have the possibility to be used as treatment for GC and COVID-19.

**Table 4.**
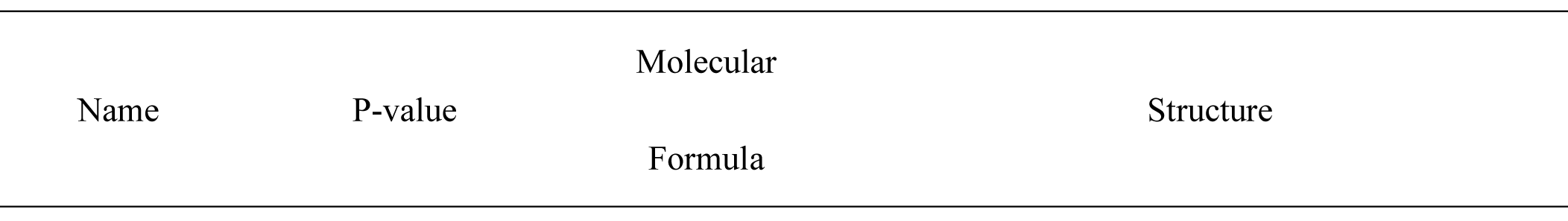
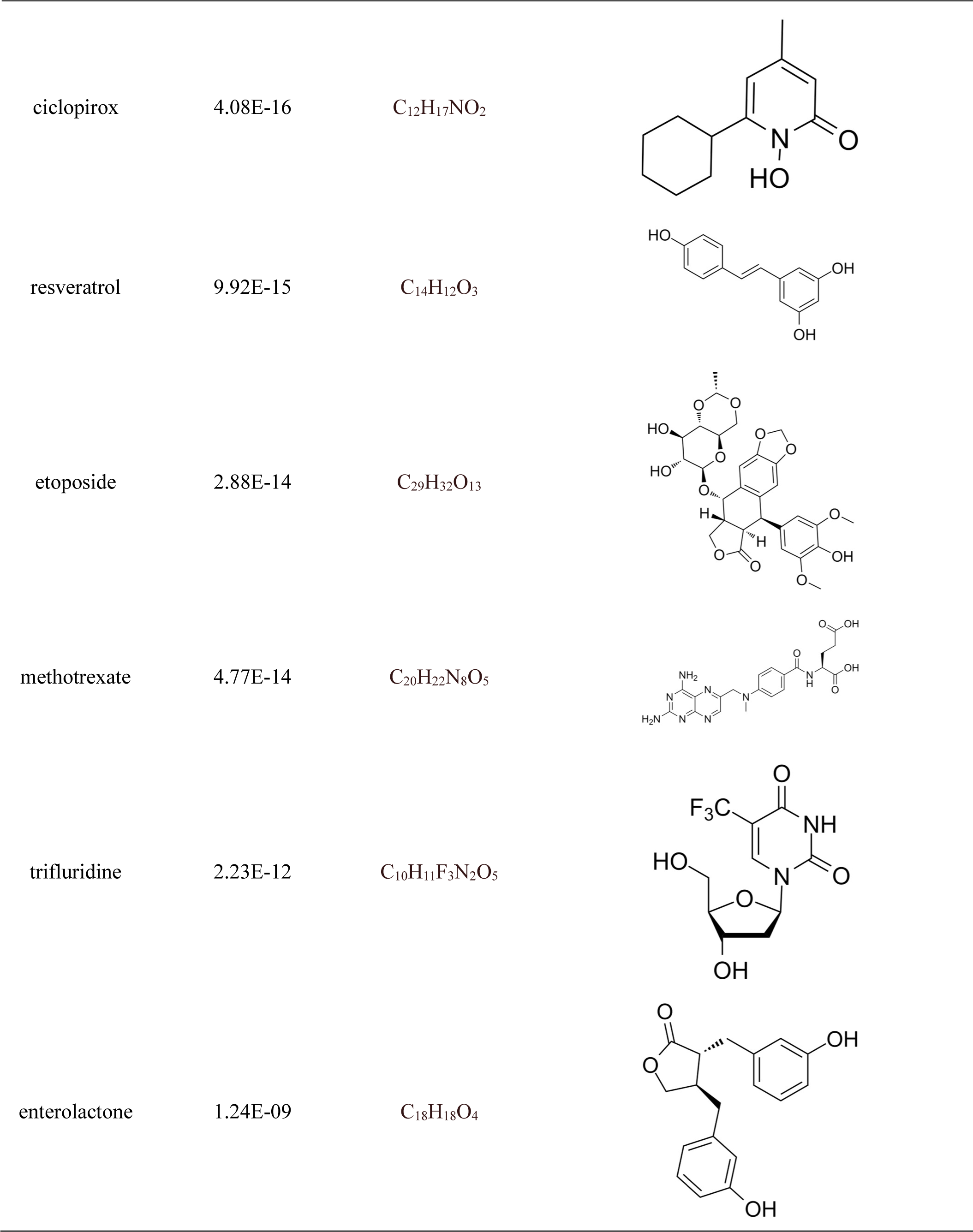
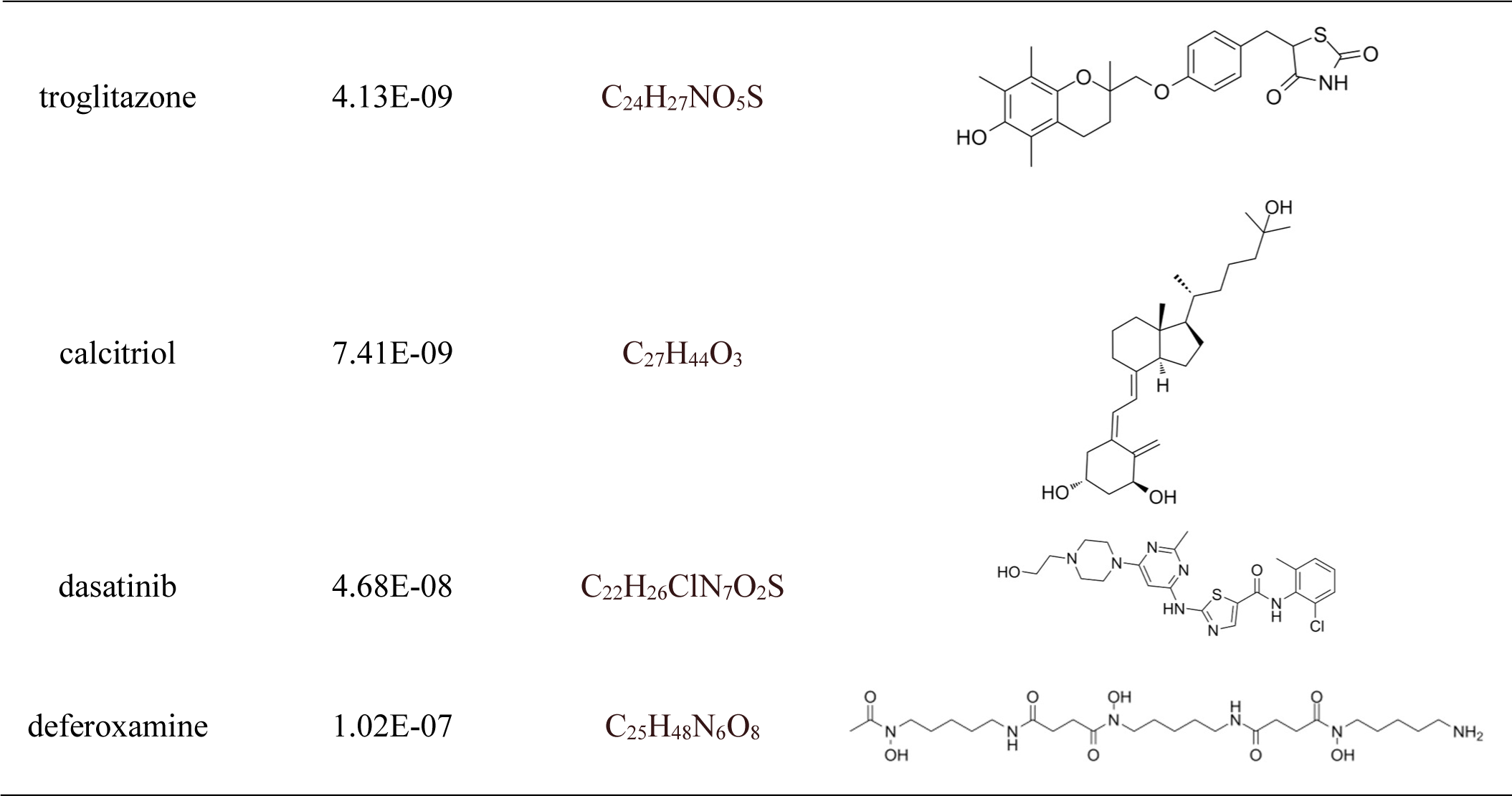
Drug candidates combined with hub genes

| Name | P-value | Molecular<br>Formula | Structure |
| --- | --- | --- | --- |
| ciclopirox | 4.08E-16 | C<sub>12</sub>H<sub>17</sub>NO<sub>2</sub> |  |
| resveratrol | 9.92E-15 | C<sub>14</sub>H<sub>12</sub>O<sub>3</sub> |  |
| etoposide | 2.88E-14 | C<sub>29</sub>H<sub>32</sub>O<sub>13</sub> |  |
| methotrexate | 4.77E-14 | C<sub>20</sub>H<sub>22</sub>N<sub>8</sub>O<sub>5</sub> |  |
| trifluridine | 2.23E-12 | C<sub>10</sub>H<sub>11</sub>F<sub>3</sub>N<sub>2</sub>O<sub>5</sub> |  |
| enterolactone | 1.24E-09 | C<sub>18</sub>H<sub>18</sub>O<sub>4</sub> |  |
| troglitazone | 4.13E-09 | C<sub>24</sub>H<sub>27</sub>NO<sub>5</sub>S |  |
| calcitriol | 7.41E-09 | C<sub>27</sub>H<sub>44</sub>O<sub>3</sub> |  |
| dasatinib | 4.68E-08 | C<sub>22</sub>H<sub>26</sub>ClN<sub>7</sub>O<sub>2</sub>S |  |
| deferoxamine | 1.02E-07 | C<sub>25</sub>H<sub>48</sub>N<sub>6</sub>O<sub>8</sub> |  |

### Exploration of gene-disease associations

Different diseases can generally be considered to be associated with each other if they have one or more similar genes. With the DisGeNET database, Network-Analyst was used to analyse gene-disease associations (Figure 9). Network-Analyst further revealed the stomach neoplasms, colonic neoplasms, autosomal recessive predisposition, liver cirrhosis experimental, mammary neoplasms, neoplasm metastasis, and prostatic neoplasms to be most associated with the identified COVID-19/GC- related common DEGs.

**Figure 9.**
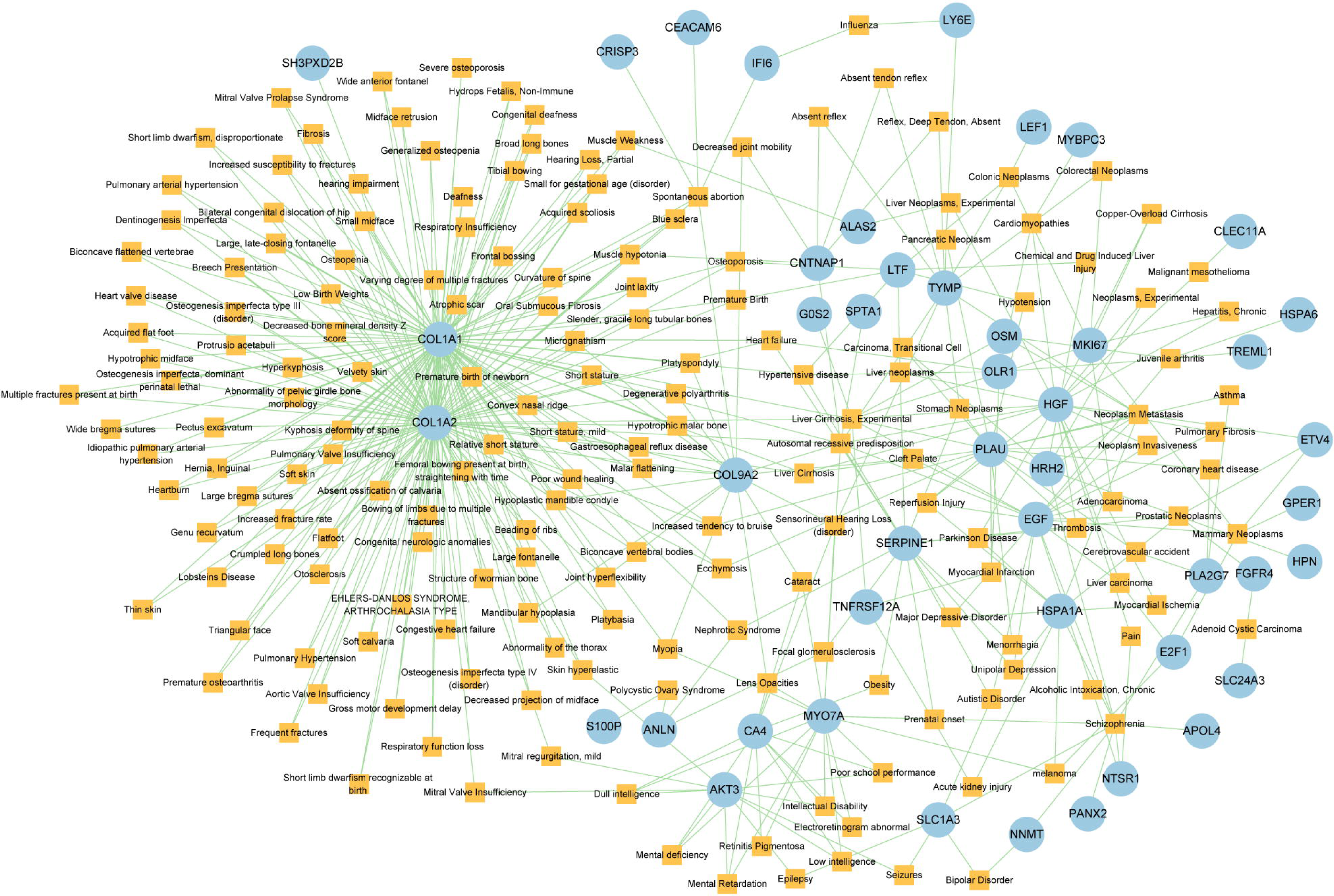
The Gene-disease association network. The square nodes represent diseases and the round nodes represent DEGs.

## Discussion

A strong correlation between COVID-19 and GC has been reported[24,26], and in addition, the receptor ACE2 of SARS-CoV-2 infected hosts is expressed in the gastric mucosa[25]. Gastric cancer patients are relatively more susceptible to viral infection than ordinary people after surgery, chemotherapy or radiotherapy. Once infected by the virus, the disease progresses relatively more quickly, has a higher severity, and has a greater risk of death. Therefore, it is important to explore the link, interaction, and co-pathogenesis between GC and COVID-19.

Here, we identified 209 common DEGs between COVID-19 and GC and explored the biological function of shared DEGs in the pathogenesis of COVID-19 and GC. Notably, these common DEGs are significantly enriched in many immune-related pathways. Alterations in neutrophil number and function have been identified as one of the immunopathological markers associated with severe COVID-19[46]. Studies have shown that neutrophils in healthy individuals can become dysfunctional in degranulation due to factors secreted by epithelial cells which were infected by SARS-CoV-2[47]. Also, neutrophils play an important role in tumour progression and metastasis[48]. Cytokine activity was affected by SARS-CoV-2 infection. Impaired acquired immune responses and uncontrolled inflammatory innate responses to SARS-CoV-2 may lead to cytokine storms[49]. The occurrence and development of tumours are actually closely related to the immune system, and cancer patients generally have immune dysfunction and low resistance. At the same time, in the process of tumour treatment, including surgery, chemotherapy and radiotherapy, it has a great impact on the body, which will have a certain impact on the patient’s immune system.

The common DEGs are utilized to construct the PPI network, in which the hub gene is the most significant regulator in the common pathogenetic processes of GC and COVID-19. CDK1 is an important regulator of cell cycle at G1/S and G2/M checkpoints[50]. It has been reported that CDK1 is highly expressed in gastric cancer. Phosphorylation of islet-1 serine 269 by CDK1 can increase its transcriptional activity and promote the proliferation of gastric cancer cells[51], and inhibition of CDK1 can inhibit the proliferation, migration and invasion of GC cells[52]. CDK1 is highly expressed in PBMCs of COVID-19 patients and is involved in the apoptosis process; CDK1 may also be associated with a worsening of the course of COVID-19, which is characterized by an extreme decrease in immune cells[53]. KIF20A (also known as mitotic kinesin-like protein 2, MKlp2) transports chromosomes during mitosis and plays a key role in cell division. KIF20A is highly expressed in almost all cancers, including gastric cancer[54], melanoma[55], hepatocellular carcinoma[56], and breast cancer[57]. Several studies have also shown that KIF20A is a hub gene involved in SARS-coV-2 infection[58,59]. TPX2 is a microtubule-associated protein that activates the cell cycle kinase protein Aurora-A, which then plays an vital role in spindle formation in mitosis[60], and high TPX2 expression is associated with tumor progression and low survival in gastric cancer[61]. TPX2 may be a novel COVID-19 intervention target and biomarker[62]. Overexpression of UBE2C is associated with poor prognosis of patients with gastric cancer, and it is also a potential biomarker for intestinal-type gastric cancer[63]. A study of peripheral blood transcriptome sequencing in patients with pneumonia found that the expression of UBE2C in patients with severe pneumonia was higher than that in patients with mild pneumonia[64]. HJURP[65], CENPA[66], PLK1[67] and IFI6[68] were found to significantly increase in gastric cancer tissues compared with normal tissues. In addition, HJURP[69], PLK1[58], MKI67[70] and IFI6[71] have been identified as potential therapeutic target for COVID-19 patients. IFIT2 have been proved to have important roles in regulating apoptosis. Chen et al. showed that decreased expression of IFIT2 promotes gastric cancer progression and predicts poor patient prognosis[72]. Similarly, the significant downregulation of IFIT2 has been observed in patients with severe COVID-19[73].

In this study, we identified a variety of compounds and medications that may treat COVID-19 and GC, including ciclopirox, resveratrol, etoposide, methotrexate, trifluridine, enterolactone, troglitazone, calcitriol, dasatinib and deferoxamine. Ciclopirox is an antifungal drug that was recently identified as a promising cancer treatment[74]. Ciclopirox regulates the growth and autophagic cell death of GC cells by regulating the phosphorylation of STAT3 at Tyr705 and Ser727 residues, and suggests that ciclopirox may be a potential treatment for GC[75]. Consistent with this study, Zhang et al. identified ciclopirox as a potential therapeutic agent for the treatment of patients with SARS-CoV-2 infection through drug prediction and simulated docking patterns[76]. Resveratrol is considered an anti-inflammatory and antiviral agent. Resveratrol inhibits the progression of gastric cancer by anti-inflammatory, antioxidant[77], antibacterial[78], inducing cell cycle arrest[79], promoting apoptosis[80], and inhibiting proliferation[81]. Besides, resveratrol downregulates neutrophil extracellular traps (NETs) generation by neutrophils in patients with severe COVID-19[82]. Similarly, network pharmacology has shown that resveratrol can alleviate COVID-19-related hyperinflammation[83]. Etoposide is a class of anticancer drugs[84]. Etoposide induces cell death through the mitochondria-dependent effects of p53[85]. The interaction of etoposide with pertuzumab or trastuzumab induces programmed cell death in gastric cancer cells through exogenous and endogenous apoptotic pathways[86]. Besides, etoposide may be a very effective treatment to protect critically ill patients from death caused by a storm of COVID-19-specific cytokines[87]. Methotrexate is a tightly bound dihydrofolate reductase (DHFR) inhibitor that is used both as an antineoplastic agent and as an immunosuppressant[88]. Clinical and experimental data suggest that methotrexate has a protective effect on SARS-CoV-2 infection by downregulating ACE2[89]. Trifluridine/tipiracil could be a new treatment option for patients with heavily pretreated advanced gastric cancer after progression on, or intolerance to, two or more previous lines of chemotherapy, including a fluoropyrimidine, a platinum agent, a taxane or irinotecan (or both), and an anti-HER2 therapy (in patients with HER2-positive disease)[90]. Similarly, trifluridine is considered to have good anti-SARS-CoV-2 capability by virtual screening, ADME/T, and binding free energy analysis[91,92]. Enterolactone is a bioactive phenolic metabolite known as mammalian lignans derived from dietary lignans[93]. Enterolactone has potent anti-cancer and/or protective properties against different cancers, including gastric[94], breast[95], colorectal[96], lung[97], ovarian, endometrial[98], and hepatocellular carcinoma[99]. In another bioinformatics and systems biology analysis, enterolactone was similarly identified as a potential treatment for COVID-19[70]. Troglitazone induces apoptosis in gastric cancer cells through the NAG-1 pathway[100]. In addition, troglitazone has been identified as a potential inhibitor of SARS-CoV-2 replicase[101]. Calcitriol alleviates COVID-19 complications by modulating pro-inflammatory cytokines, antiviral proteins, and autophagy[102]. Similarly, there was an improvement in peripheral arterial oxygen saturation and inspired oxygen fraction in hospitalized patients with COVID-19 treated with calcitriol[103]. Dasatinib promotes TRAIL-mediated apoptosis by upregulating CHOP-dependent death receptor 5 in gastric cancer[104]. Dasatinib can reduce SARS-CoV-2-related mortality, delay its onset, and reduce the number of other clinical symptoms[105]. Deferoxamine is a widely used iron chelator used to treat iron overload. Deferoxamine targets mitochondria and impair mitochondrial respiration and [Fe-S] cluster/heme biogenesis in cancer cells, thereby inhibiting tumor proliferation and migration and inducing cell death[106]. Deferoxamine has iron chelation, antiviral, and immunomodulatory effects to help control SARS-CoV-2[107].

## Conclusions

In this study, transcriptome analysis was applied to summarize the relationship between gastric cancer and COVID-19. DEGs for GC and COVID-19 were obtained in the GEO dataset, 209 shared DEGs were identified, and associations between gastric cancer and COVID-19 were found. To clarify what role these DEGs play at the transcriptional level, enrichment analysis was conducted. We also used these common DEGs to obtain a PPI network and defined the 10 most important hub genes: *CDK1*, *KIF20A*, *TPX2*, *UBE2C*, *HJURP*, *CENPA*, *PLK1*, *MKI67*, *IFI6*, and *IFIT2*. Besides, we established a TF-gene and miRNA-gene interaction network for hub genes and identified key TFs and miRNAs. More importantly, we identified a variety of compounds and drugs that may treat COVID-19 and GC, such as ciclopirox, resveratrol, etoposide, methotrexate, trifluridine, enterolactone, troglitazone, calcitriol, dasatinib and deferoxamine. This study shows new possibilities for the treatment of COVID-19 and GC.

## Acknowledgements

We are most grateful for GEO databases providing their platform and thanking contributors who have uploaded their valuable datasets. Additionally, we appreciated Core Facility of West China Hospital for their technique support. Also, the graphic abstract of the overall general workflow for this study was created with BioRender.com

## Author contribution

Zhen Wang and Kefei Yuan were responsible for funding acquisition. Tengda Huang was responsible for design conception. Hongyuan Pan and Yong Zeng were responsible for formal analyses. Xiao Ma performed data curation. Xinyi Zhou and Ao Du verified the underlying data. Tengda Huang and Xiaoquan Li prepared the original manuscript draft. All authors contributed to manuscript reviewing and editing. All authors contributed to the article and approved the submitted version.

## Data availability statement

The datasets analyzed during the current study are available in the GEO database, https://www.ncbi.nlm.nih.gov/geo/query/acc.cgi?acc=GSE196822 and https://www.ncbi.nlm.nih.gov/geo/query/acc.cgi?acc= GSE179252.

## Consent for publication

Not applicable.

---

**Table S1** The differentially expressed genes of COVID-19.

**Table S2** The differentially expressed genes of GC.

**Table S3** Common DEGs between COVID-19 and GC.

**Table S4** Hub genes in network string_interactions ranked by MCC method.

**Table S5** The TFs and their interactions with hub genes.

**Table S6** The microRNAs and their interactions with hub genes.

